# In Silico transcriptional analysis of asymptomatic and severe COVID-19 patients reveals the susceptibility of severe patients to other comorbidities and non-viral pathological conditions

**DOI:** 10.1101/2022.04.16.488556

**Authors:** Poonam Sen, Harpreet Kaur

## Abstract

COVID-19 is a severe respiratory disease caused by SARS-CoV-2, a novel human coronavirus. The host response to SARS-CoV-2 infection is not clearly understood. Patients infected with SARS-CoV-2 exhibit heterogeneous intensity of symptoms, i.e., asymptomatic, mild, and severe. Moreover, effects on organs also vary from person to person. These heterogeneous responses pose pragmatic hurdles for implementing appropriate therapy and management of COVID-19 patients. Post-COVID complications pose another major challenge in managing the health of these patients. Thus, understanding the impact of disease severity at the molecular level is vital to delineate the precise host response and management. In the current study, we performed a comprehensive transcriptomics analysis of publicly available seven asymptomatic and eight severe COVID-19 patients. Exploratory data analysis using Principal Component Analysis (PCA) showed the distinct clusters of asymptomatic and severe patients. Subsequently, the differential gene expression analysis using DESeq2 identified 1,224 significantly upregulated genes (logFC>= 1.5, p-adjusted value <0.05) and 268 significantly downregulated genes (logFC<= -1.5, p-adjusted value <0.05) in severe samples in comparison to asymptomatic samples. Eventually, Gene Set Enrichment Analysis (GSEA) of upregulated genes revealed significant enrichment of terms, i.e., anti-viral and anti-inflammatory pathways, secondary infections, Iron homeostasis, anemia, cardiac-related, etc. Gene set enrichment analysis of downregulated genes indicates lipid metabolism, adaptive immune response, translation, recurrent respiratory infections, heme-biosynthetic pathways, etc. In summary, severe COVID-19 patients are more susceptible to other health issues/concerns, non-viral pathogenic infections, atherosclerosis, autoinflammatory diseases, anemia, male infertility, etc. And eventually, these findings provide insight into the precise therapeutic management of severe COVID-19 patients and efficient disease management.

## 1. Introduction

Since its first reported case at the end of 2019, an acute respiratory syndrome causing novel Coronavirus (2019-nCoV) outbreak in the human population has taken the world by storm. The 2019-nCoV was later officially named SARS-CoV-2 (**S**evere **A**cute **R**espiratory **S**yndrome related novel **Co**rona**v**irus **2**), and the disease caused by it COVID-19 (**Co**rona**vi**rus **D**isease 2019) [1–3]. The virus spread uncontrollably so much that in January 2020, World Health Organization declared COVID-19 as a “public health emergency of international concern” (PHEIC) and eventually as a pandemic in March 2020 [1]. As of 1^st^ April 2022, the total reported cases worldwide stand at 488,190,137 [4]. The SARS-CoV-2 is an enveloped, positive single-stranded RNA virus that belongs to the Coronaviridae family, β-coronavirus genus, and is believed to have a zoonotic to human transmission [3, 5, 6]. The trimeric spike (S) protein that forms the virus’s envelope plays an essential role in the virus-host cell interaction [7]. There are six other coronaviruses, i.e., 229E, OC43, NL63, HKU1, SARS-CoV, and MERS-CoV, which are already known to infect humans and cause respiratory and gastrointestinal problems [8]. These human coronaviruses (HCoVs) are generally considered inconsequential except for our experience with SARS-CoV in 2003, MERS in 2012, and SARS-CoV-2 with the ongoing pandemic [9].

The mutations in the viral spike protein components, especially in its receptor-binding domain, have resulted in the generation of multiple variants, of which Delta variant (B.1.617.2) became a “variant of concern” (VOC) and posed a significant threat to human health [10–12]. Our health sector has faced major challenges in tackling disease spread and providing management of symptoms in the patients [13]. Multiple drugs are introduced for symptomatic treatments, but none has been efficient to treat all symptoms caused due to the viral infection [14]. Even a few drugs that were believed to be helpful in COVID-19 disease management were later found to cause other health concerns in the patients administered with these [15]. The difficulty faced in devising standard therapeutic options is due to the high mutability rate of the virus, a complex interplay of virus-host interaction, and an individual’s immune response to the infection [16–19].

SARS-CoV-2 impacts individuals in peculiar ways [16]. Most infected subjects are asymptomatic or mildly symptomatic, but some develop severe symptoms [16]. Comorbidities such as diabetes mellitus, hypertension, cardiovascular disease (CVD), and advanced age further increase the risk of disease severity [20–23]. As in many asymptomatic or mild cases, diagnostic test reports false-negative results even in the presence of infection, and due to the shared spectrum of symptoms with other viral infections, it becomes difficult to discern COVID-19 from other viral infections [24]. This makes disease management further complicated. Another primary concern associated with COVID-19 is high infectivity as it spreads by human contact and through air droplets and aerosols, making it difficult to control [25, 26]. COVID-19 spread through fecal matter is speculated in some studies, though the presence of viral particles in the fecal samples of infected individuals is well documented and makes it an essential diagnostic tool [27, 28]. The main clinical manifestations of SARS-CoV-2 in severe COVID-19 patients involve lower respiratory tract issues resulting in Acute Respiratory Distress Syndrome (ARDS) and hypoxia, fever, cytokine storm due to hyperactive immune system, brain fog, headache, cardiac arrest, and muti-organ damage and even death in severe cases [22, 29–33]. Most disease symptoms may persist for 10-15 days, with some may exist for a prolonged time [34, 35]. It is well known that even after the viral load declines significantly, many health issues persist in the COVID-19 recovered patients [36–38]. These post-COVID effects are observed mainly in hospitalized and severe patients and add to another layer of disease mismanagement [39, 40]. So, the significant challenges of disease management include SARS-CoV-2’s high infectivity rate, poor efficacy of available treatments, the complexity of symptoms, and less understanding of disease progression [41]. SARS-CoV-2, upon entry into the nasopharyngeal tract, interacts with the transmembrane serine protease 2 (TMPRSS2) and Angiotensin-Converting Enzyme 2 (ACE2) receptors present on the endothelial cells of the respiratory tract [42]. ACE2 receptors are also present in other organs, such as the gastrointestinal tract, lymph nodes, thymus, bone marrow, spleen, liver, kidney, skin, and brain. This might be the possible reason for the viral impact on these organs [33, 43–46]. As extensively studied, virus entry in these organs is mediated through the interaction of receptor-binding domain on spike protein of virus and the ACE2 receptors present on host cells [45, 47]. Upon infection, the virus replicates inside the host cell using the host replication machinery. In response to all this, the host immune system fights to reduce the viral load by inhibiting the replication of viral RNA. The diverse symptoms results from the involvement of various biochemical pathways triggered by viral entry and replication, the host cellular response to control the spread of the infection [48].

With the advancement in the RNA sequencing technology, one can view the transcriptomic landscape under a given condition and for a particular cell type. It is also instrumental in understanding the pathogenesis of a disease in the host [49]. Diverse scientific groups across the globe have developed numerous resources and tools to compile and analyze the data from host and pathogen [50–73].

Due to the systemic effects of COVID-19 infection, it becomes more challenging to treat patients with complex symptoms. Hence, we believe studying the differential mechanism operating in asymptomatic and severe COVID-19 patients can help us manage disease manifestations in severe patients. Thus, we performed a comparative analysis of transcriptomics profiles of severe and asymptomatic COVID-19 patients using PCA and DESeq2. Exploratory analysis based on PCA of samples shows two clear, distinct clusters of severe and asymptomatic samples. The differential gene expression analysis revealed significantly altered transcriptomics patterns between these two groups. Subsequently, Gene Set Enrichment Analysis (GSEA) identified some of the key altered pathways and biological processes involved in severe patients compared to asymptomatic patients.

## 2. Methods

### 2.1. Dataset and Experimental design

In the current study, we obtained publicly available data (GSE178967) from the NCBI GEO (Gene Expression Omnibus). This dataset comprises RNA-Seq read counts and metadata information conducted on 108 SARS-CoV-2 subjects by the Stanford COVID-19 CTRU [74]. These COVID-19 subjects, confirmed by RT-PCR, were administered Peginterferon Lambda and placebo on day00. Peginterferon Lambda is a therapeutic drug for reducing the viral particles in COVID-19 patients [75]. Whole blood samples for RNA extraction for high throughput sequencing were collected on day 00 (untreated) and day 05 (treated) from the day of drug administration. The available RNA sequencing data are the read counts aligned to transcripts or genes for 180 samples from day 00 and day 05 of 108 subjects. In the series matrix file (provided in GEO), the COVID-19 subjects are categorized as asymptomatic, moderately symptomatic, and severe [74]. We have also used the same categorization of subjects for our analysis. The series matrix file contains other clinically significant information such as age, gender, day from drug administration (Peginterferon Lambda and placebo), and viral shedding value. The details and data structure of the study are summarized in Table 1 and Supplementary Table S1, respectively. The summary of clinical information extracted from the GEO series matrix is provided in Supplementary Table S2.

**Table 1:** Detail of the study as derived from GEO [1].

|  |  |
| --- | --- |
| **GEO accession** | Series GSE178967 |
| **Study title** | Baseline signatures associated with clinical, virologic, and immunologic outcomes in patients with mild to moderate COVID-19 |
| **Organism** | *Homo Sapiens* |
| **Bio project** | PRJNA741686 |
| **SRA** | SRP325729 |
| **Platform** | GPL24676<br>Illumina NovaSeq 6000 |

### 2.2. Data Preparation and normalization

#### 2.2.1. Data Pre-processing

The data contains sample IDs in row 1, transcript IDs (ENST ID) in column 1, gene symbols in column 2, and corresponding non-normalized read counts in the matrix as integer values. The sample IDs belong to asymptomatic, moderately symptomatic, or severe subjects from day 0 or day 5 of peginterferon lambda and placebo administration. The RNA sequencing expression values of the dataset are non-normalized read counts (as mentioned in supplementary file information of the original dataset submitted in GEO) [74]. These read counts are the number of reads mapped and aligned to a particular transcript/gene region identified from the human reference genome. It is generally required to pre-process the read count data to get statistically significant results [76–80]. We followed common pre-processing steps for both PCA and Differential Gene Expression (DGE) analysis, but the normalization steps were different based on the downstream analyses. The Principal Component Analysis is a dimensionality reduction unsupervised machine learning method that requires normalized data [78, 79, 81, 82]. While DESeq2 is a DGE analysis tool that mandates data to be unprocessed read counts as integer values [83]. DESeq2 uses inbuilt methods to normalize for library size and hence does not require prior normalization [83–85].

The summary of workflow, including pre-processing and normalization, is depicted in Figure 1. In pre-processing, we removed rows with NA, taken the average of duplicates genes using aggregate function in R, and filtered out the genes having zero or low expression. Studies suggest that low expression genes negatively impact the Differentially Expressed Gene analysis [86]. Thus, genes with low or zero variance filtered out using the nzv (non-zero variance) function of the “Caret package” available in R [87]. Genes with zero variance across all samples are considered insignificant as these do not contribute to statistical significance and only increase time in the analysis [77, 88]. After removing genes with low expression values, we performed further pre-processing specific to PCA, and for DESeq2, we continued with the pre-processed and non-normalized data. Notably, we performed exploratory data analysis using PCA on normalized data while differential gene expression analysis using DESeq2 on raw read count values.

**Figure 1.**
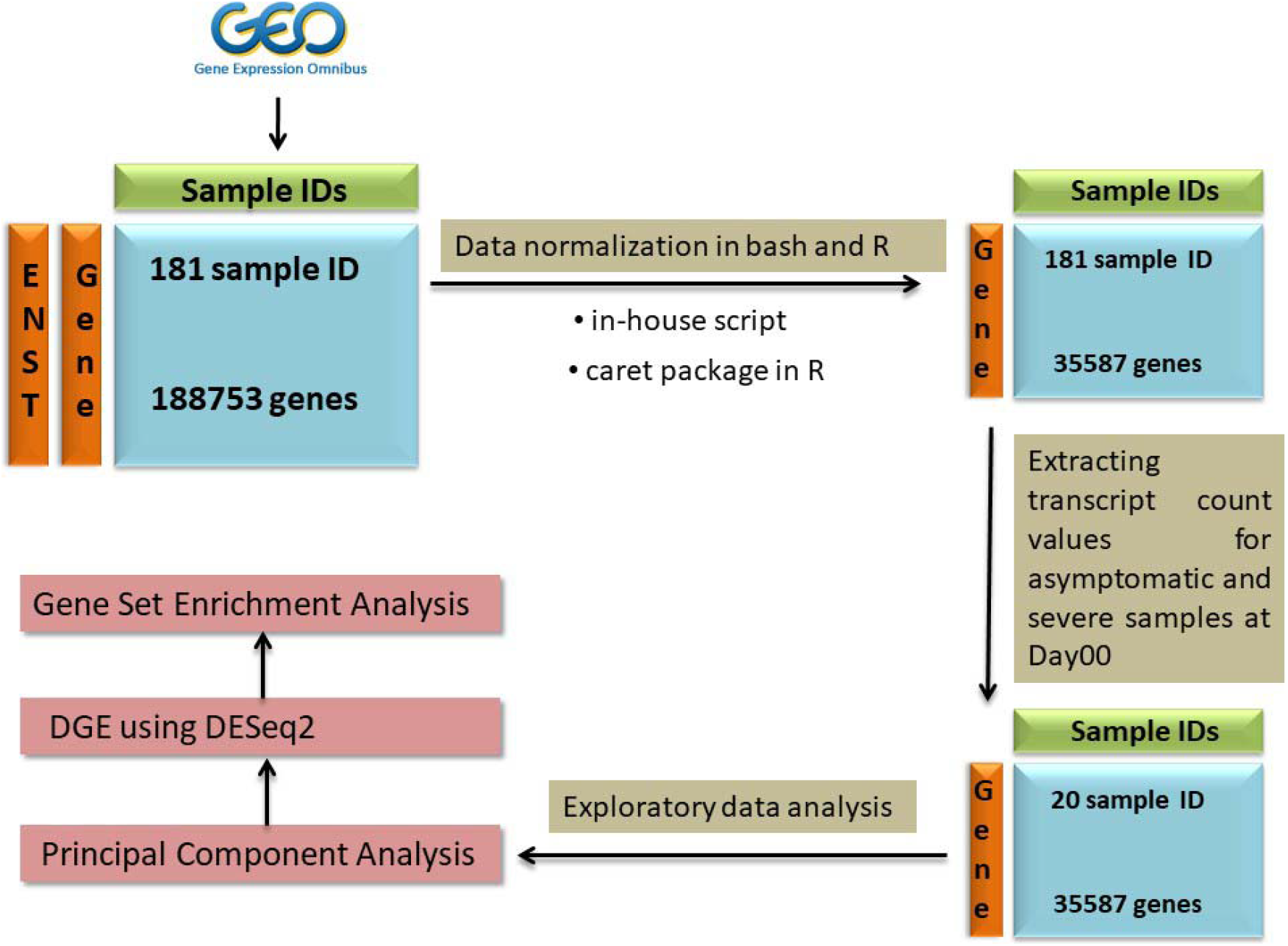
The Complete workflow of the study.

#### 2.2.2. Data Normalization

After the abovementioned pre-processing, subsequently, for PCA, we normalized the read counts by transforming them to log values and then performing center and scaling using the “Caret package” available in R [87]. The data matrix that resulted from PCA normalization contains 180 samples with log-transformed read counts for 35,587 gene rows.

### 2.3. Analysis Methodology

#### 2.3.1. Exploratory Data Analysis using Principal Component Analysis

We performed Principal Component Analysis (PCA) to identify the patterns in the dataset and variations between the samples in a group. PCA reduces the dimensions of a large dataset while retaining most of the variations. Hence, PCA assists in identifying sample clusters in a particular group and outliers [89]. We performed PCA on normalized data (comprising 180 samples with log-transformed read counts for 35,587 gene rows) using the ggfortify package in R. The first PCA includes all 180 samples of asymptomatic, moderately symptomatic, and severe subjects. Then we performed PCA for various groups as mentioned in Supplementary Table S3. One of these PCA, consists of asymptomatic and severe patients at Day 00 (untreated) which we believe will help us understand the host response mechanism in severe patients in comparison to asymptomatic. The total number of samples belonging to this group was 15, with seven asymptomatic and eight severe samples. With the help of scatterplots based on PCA components, we identified outliers, which were subsequently removed from the data for the downstream PCA and DGE analysis on untreated (Day 00) group.

#### 2.3.2. Differential gene expression analysis

After outliers removal using PCA, we performed differential gene expression analysis between severe and asymptomatic patients’ samples using the DESeq2 package in R [83]. Notably, we considered only those genes as significantly expressed between groups with a p-adjusted value <0.05. This criterion of p-adjusted value is used in numerous studies [83, 90–98]. Further, we applied another filter, i.e., Log2 fold change (Log2FC) to identify significantly upregulated (Log2FC >=1.5) and downregulated (Log2FC <=-1.5) genes in the severe patients in comparison to asymptomatic patients. Additionally, to understand patterns in gene expression between asymptomatic and severe patients, we constructed heatmaps using the heatmap function in R [99]. Heatmap is a grid-like graphical representation of the expression of genes (in rows) in all the samples (in columns) taken into consideration [100].

#### 2.3.3. Biological annotation

Subsequently, to understand the biological implication of significantly differentially expression genes obtained from DESeq2 analysis in severe patients, we performed gene enrichment analysis using the Enrichr [101–103]. We queried the upregulated and downregulated gene sets independently in the Enrichr search engine [104]. Enrichr gives various Gene Set Enrichment terms as output which can be analyzed for significance based on four ranking parameters, i.e., p-value, adjusted p-value, odds ratio, combined scores [103].

Enrichr visualization bar graph shows top enrichment terms with significance depicted by the length and color of the bar. An enrichment term with a more extended bar and a lighter shade of red indicate higher significance than a term with a shorter bar and darker red color or grey color [103]. A few of the top Gene Set Enrichment terms are, i.e., KEGG Human, WikiPathway, Gene Ontology (GO) terms, Jensen diseases, Human phenotype ontology, etc., based on p-value (<0.05). To identify significant pathways involved in each enrichment term, we used the q-value (adjusted p-value) < 0.05. Besides, we searched for the top significant and differentially expressed genes (from our analysis) in the literature to understand their already known role in COVID-19 pathogenesis.

## 3. Results

In the current study, we analyzed the transcriptomic profiles of asymptomatic and severe COVID-19 patients to compare the transcriptional changes and understand the biological implications of infection. A publicly available RNA sequencing read count dataset was extracted, pre-processed, and normalized. We performed exploratory data analysis using PCA to understand variations between groups and to identify outliers. Subsequently, we performed differential gene expression analysis between these identified groups (Severe vs. Asymptomatic). Eventually, gene enrichment analysis was performed using the significantly differentially expressed gene sets to discern their biological involvement in viral immuno-pathogenesis.

### 3.1. Data Pre-processing

After the pre-processing, we are able to remove genes without identifiers, zero expression, and low variance in the data. Thus, the total number of genes reduced from 188,753 to 35,587 in the data. Subsequently, this dataset was used for exploratory and DGE analysis.

### 3.2. Exploratory Data Analysis

We analyzed each group’s scatter plot and principal components to identify if any of the top Principal Components (PC) showed significant variations. The scatter plots for all Principal Component Analysis performed are provided in Figure 2 and Supplementary Figure 1 A-C. The scatter plot in Supplementary Figure 1.A represents all three groups, i.e., untreated and treated asymptomatic, moderately symptomatic, and severe. However, three outliers can be observed at the bottom left of the plot; the clustering does not show any clear distinction between the three groups. PCA for remaining groups, i.e., severe male v/s female, severe below 45 years age v/s above 45 years age (Supplementary Figure S1 B and C also did not show any clear groups. The top principal components in these groups also did not show significant variations. Thus, we mainly shifted our focus to two critical groups, i.e., asymptomatic (untreated or day 00) and severe (untreated or day 00), since these are two contrasting viral infection conditions and interestingly, they also represent lesser within-group variation. The PCA between untreated asymptomatic (n=7) and untreated severe samples (n=8) represent nearly 61.8% variation in the data, wherein PC1 contributes 49.49%, and PC2 contributes ~ 12.39% variation (Figure 2A). Using the clustering patterns in PCA, we identified seven samples as outliers. We removed these outlier samples and then performed PCA on the remaining eight samples (four severe and four asymptomatic), that represented nearly 48.61% variation in the data, where PC1 represents 34.64%, and PC2 represents 13.97% variation. So, Scatterplots based on the PC1 and PC2 of untreated asymptomatic and severe samples show clear distinction, and we got down to 4 samples in each group (Figure 2B). For a significant DGE analysis, the minimum number of samples in each group must be three, so we considered these four samples from both groups for further downstream analysis.

**Figure 2.**
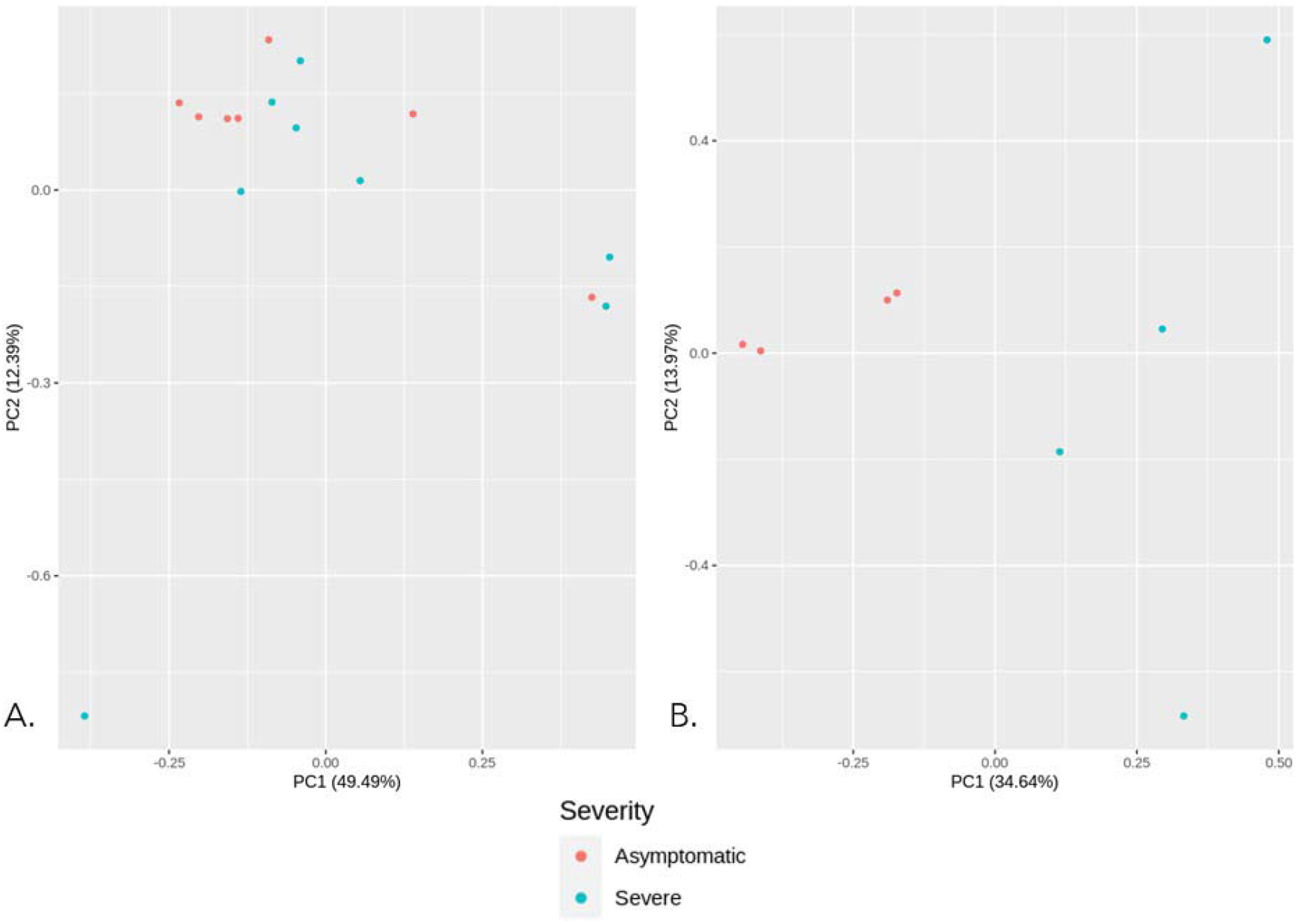
Principal Component Analysis between untreated asymptomatic and severe groups. A. PCA between asymptomatic (n=7, Day00) v/s severe samples (n=8, Day00) B. PCA between asymptomatic (n=7, Day00) v/s severe samples (n=8, Day00) after outlier removal.

### 3.3. Differential gene expression analysis

Differential gene expression analysis between untreated severe and asymptomatic samples using DESeq2 identified 2,837 genes as significantly differentially expressed (p adjusted value < 0.05). From these 2,837 genes, 1224 genes were found to be significantly upregulated (Log2FC>= 1.5, p-adjusted value <0.05) and 268 genes as significantly downregulated (Log2FC<= −1.5, p-adjusted value <0.05) in severe samples in comparison to asymptomatic samples. The list of the total up-and downregulated genes is provided in Supplementary Tables S4 and S5, respectively. The volcano plot represents the pattern of differentially expressed genes (Figure 3). Each dot in the plot represents a single gene with log2FC along the x-axis and -Log10 (p-value) along the y-axis. In the volcano plot, the genes depicted in black color are nonsignificant, while genes in blue and red color represent most significantly differentially expressed genes with padj <0.01 and padj <0.05, respectively. Further, heatmap (Figure 4) represents the expression pattern of the top 50 genes (25 upregulated and 25 downregulated genes) in untreated severe COVID-19 samples in comparison to asymptomatic samples. The color scale denotes the expression values in the heatmap. The red color’s intensity represents upregulated genes, and the yellow color’s intensity represents downregulated genes in the sample under consideration. The top 25 upregulated and down regulated genes ae mentioned in Table 2. with respective gene description.

**Figure 3.**
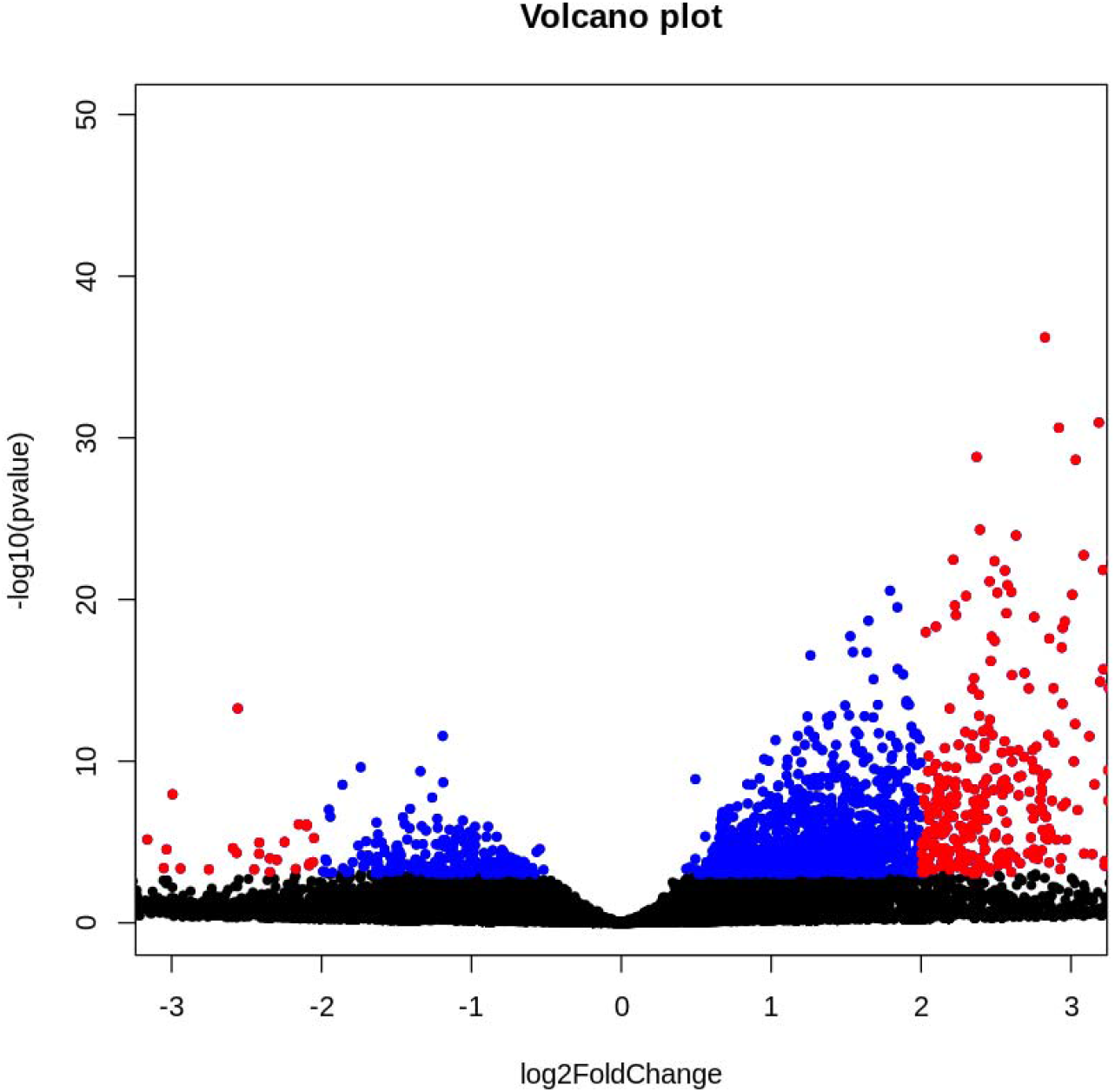
Volcano plot based on p-value and log2FC. Each dot here represents a single gene. Black represents nonsignificant genes, blue and red represent genes differentially regulated at padj <0.01 and padj <0.05.

**Figure 4.**
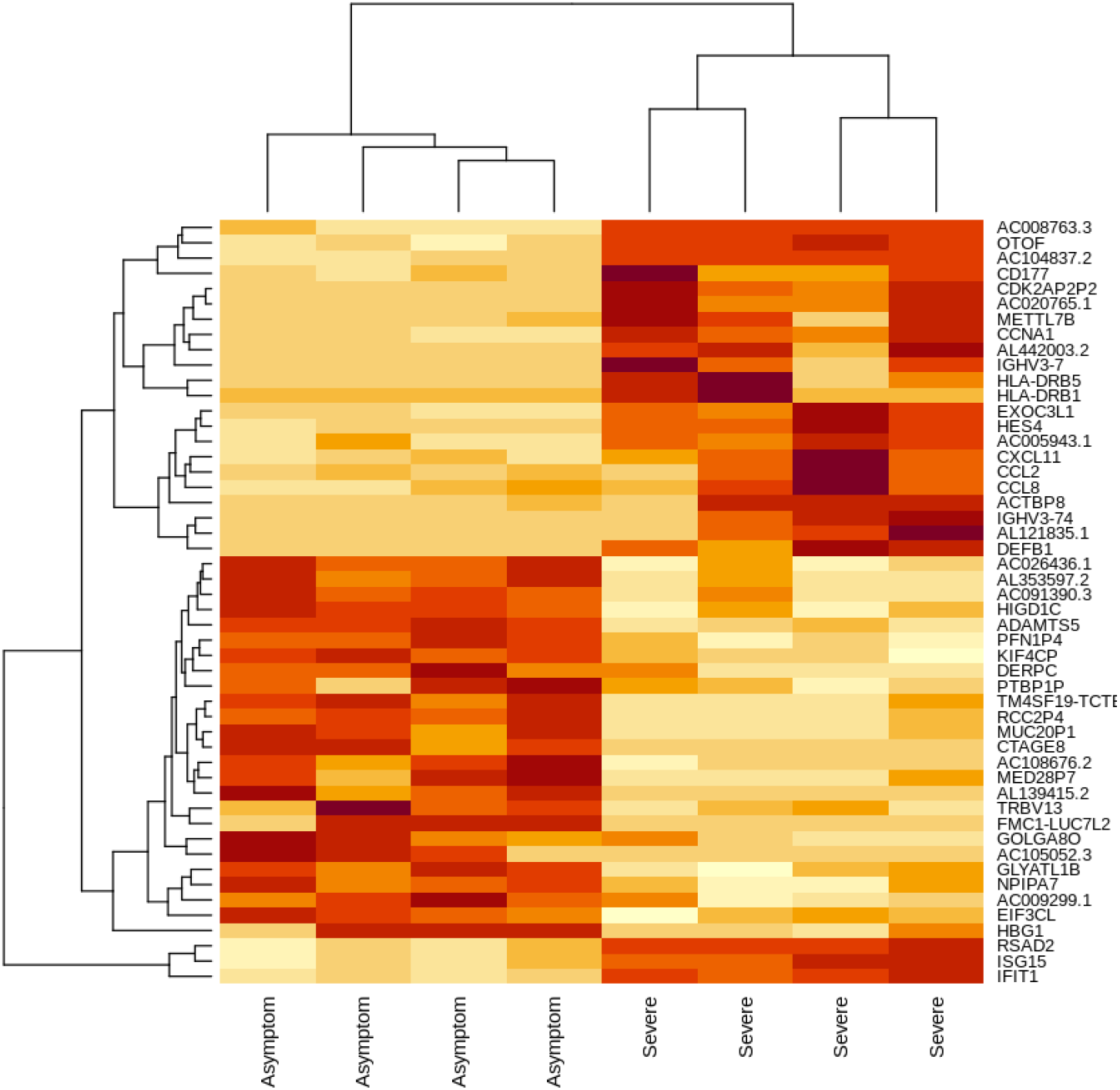
Heatmap based on top 25 upregulated and downregulated genes from DESEQ2 of asymptomatic and severe samples.

**Table 2:**
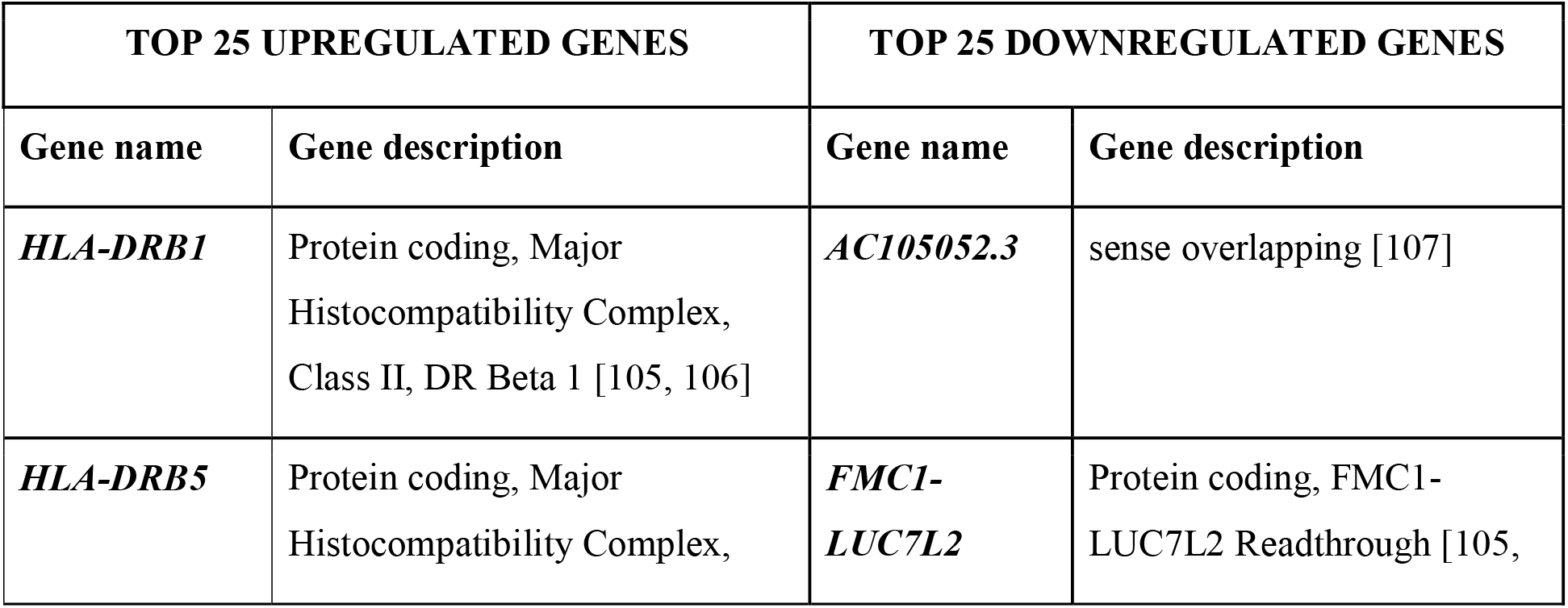
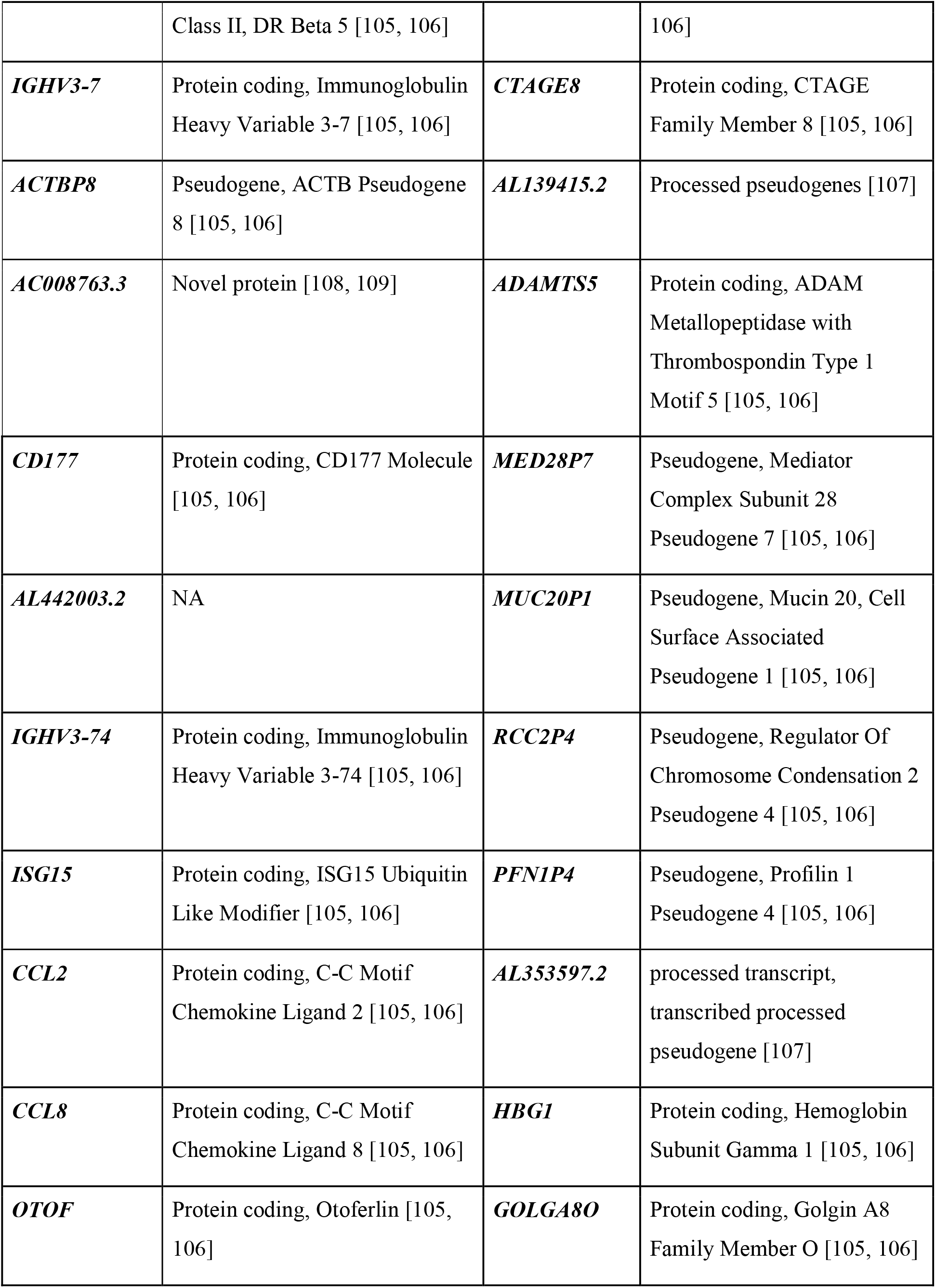
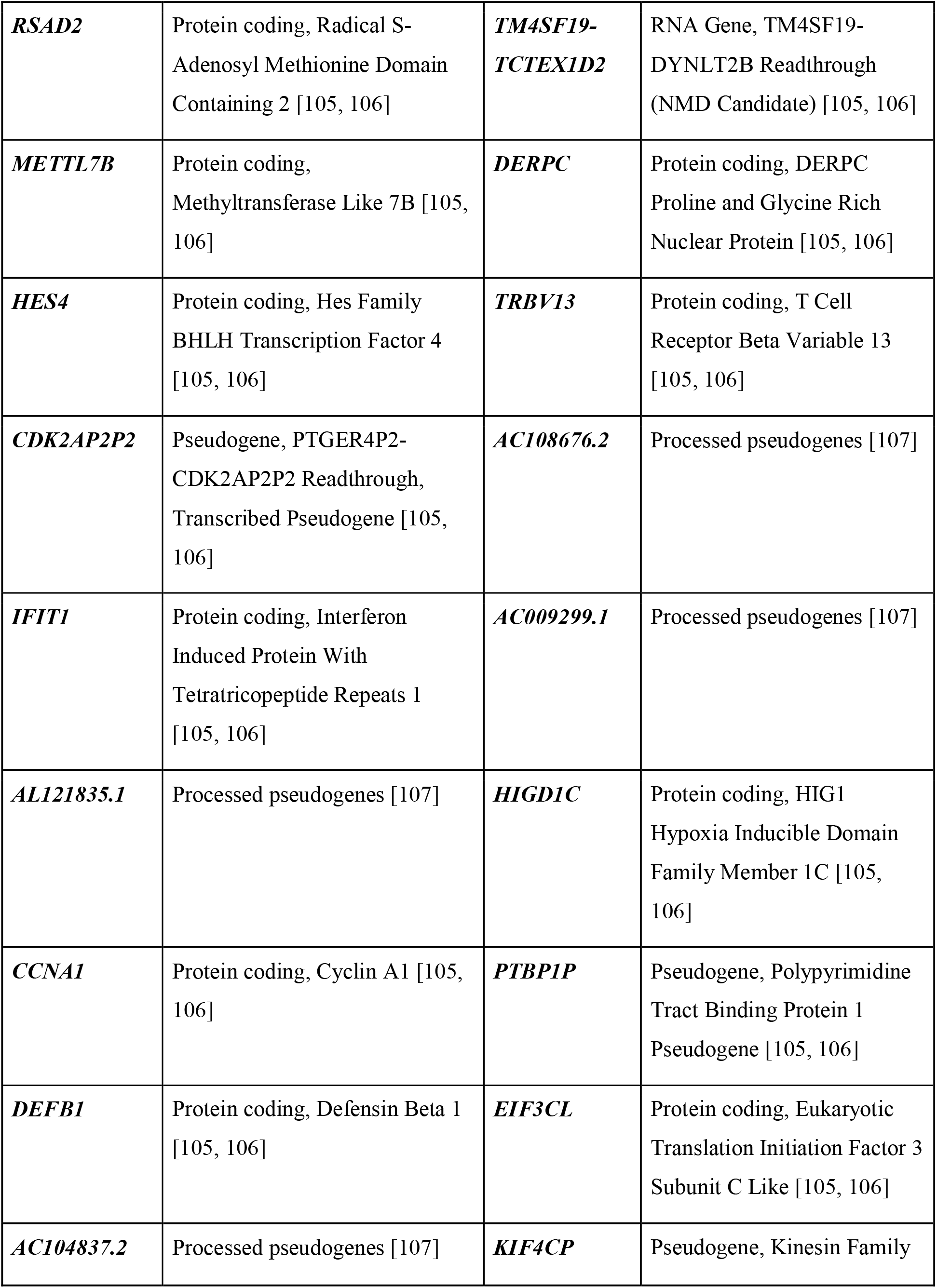
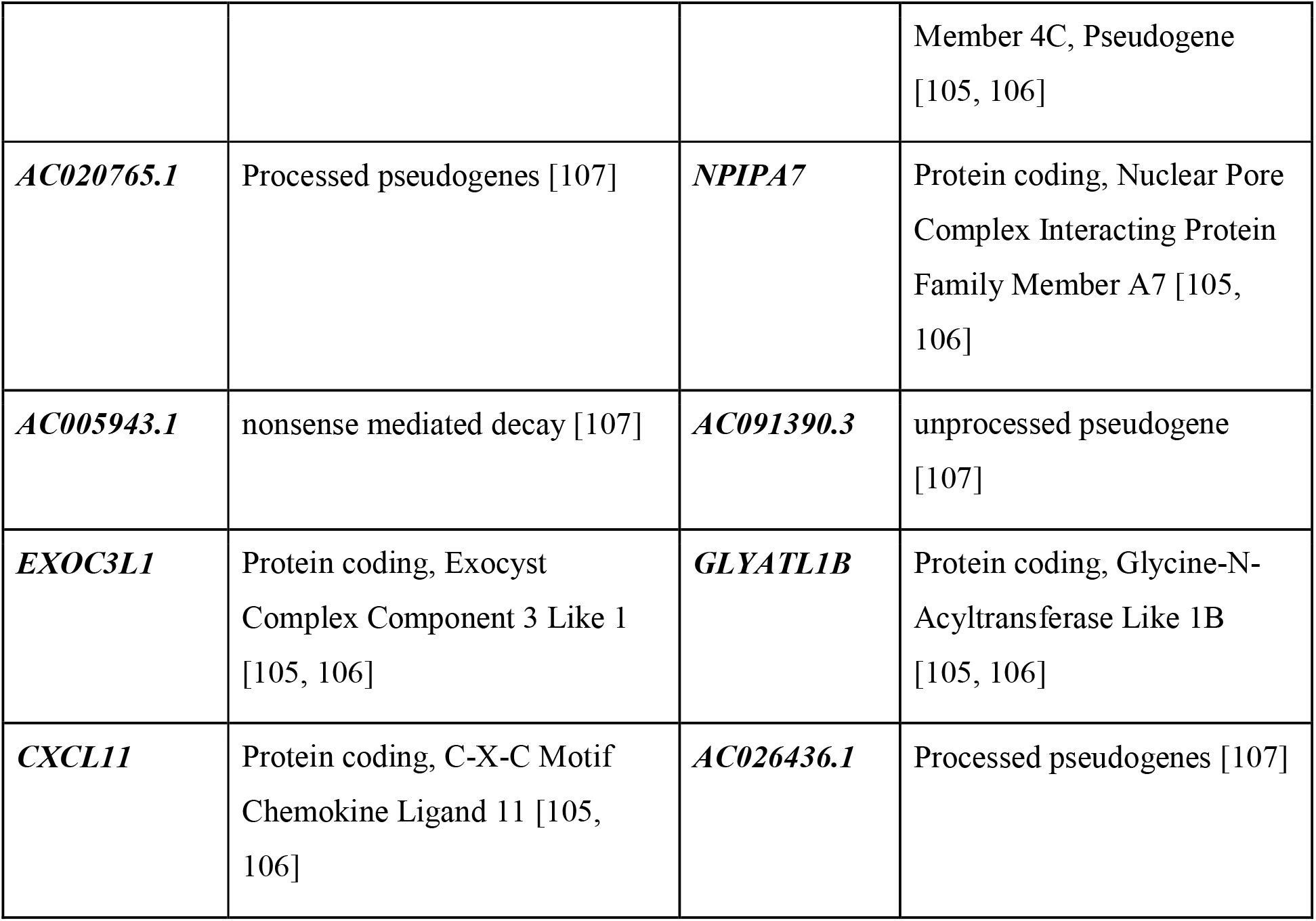
List of top 25 upregulated (Log2FC>= 1.5, p-adjusted value <0.05) and downregulated (Log2FC<= −1.5, p-adjusted value <0.05) genes in severe COVID-19 subjects in comparison to asymptomatic subjects with their gene description.

#### 3.2.5. Biological annotation - Gene Enrichment analysis

We queried all significantly up and down regulated genes obtained from DGE analysis to the Enrichr search engine independently. The resulting bar plots represent the top enriched terms for upregulated genes (Figures 5, Supplementary Figure S2-S4) and downregulated genes (Figures 6, Supplementary Figure S5). We also extracted the complete results of all enriched terms for both upregulated (see Table S6-S21, Supplementary File 2) and downregulated gene sets (see Table S22-S36, Supplementary File 2) as tables. Besides, we also studied the significant terms and searched in the literature whether these are associated with COVID-19 pathogenesis previously. The key terms are briefly explained below:

**Figure 4:**
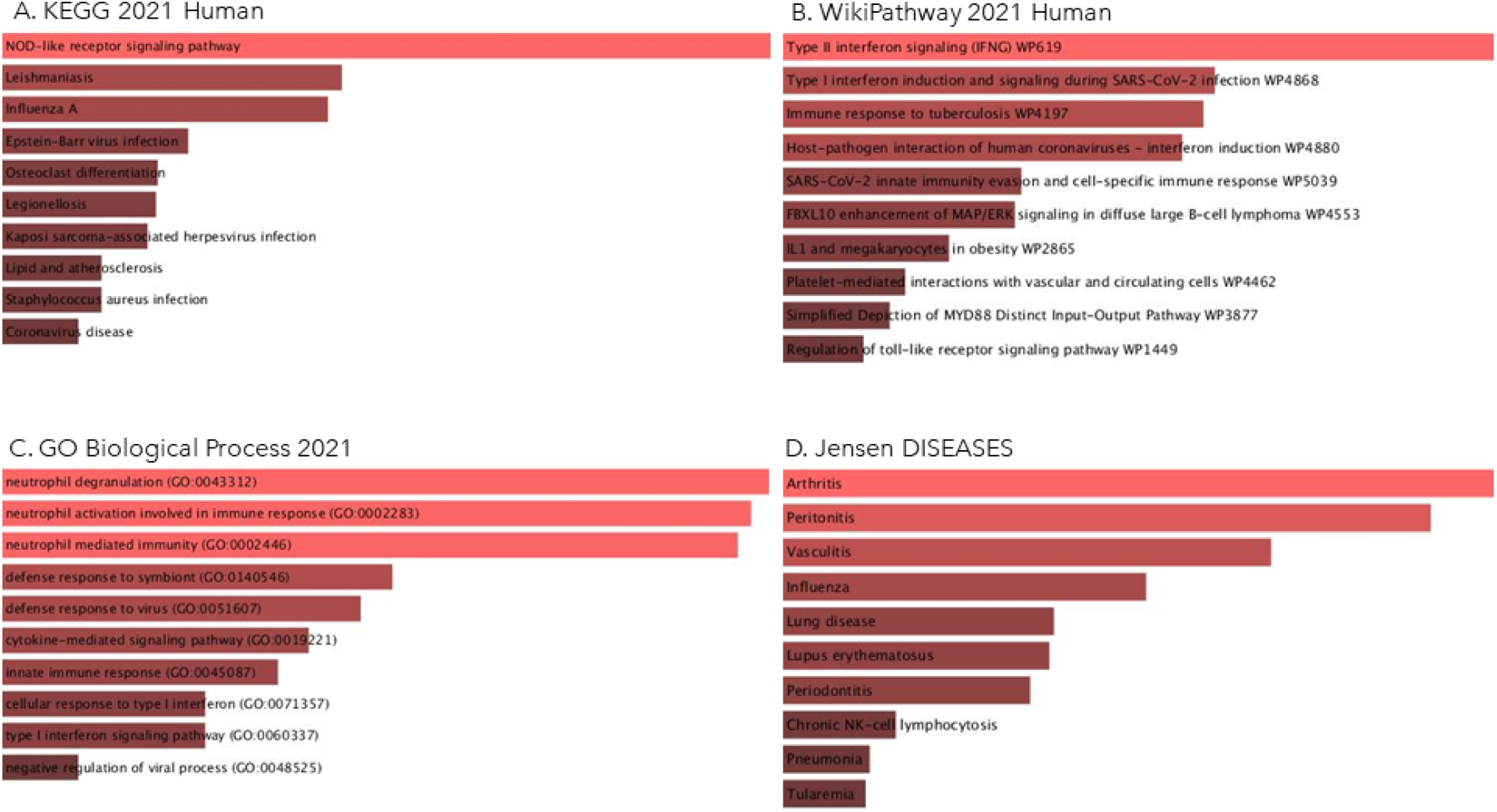
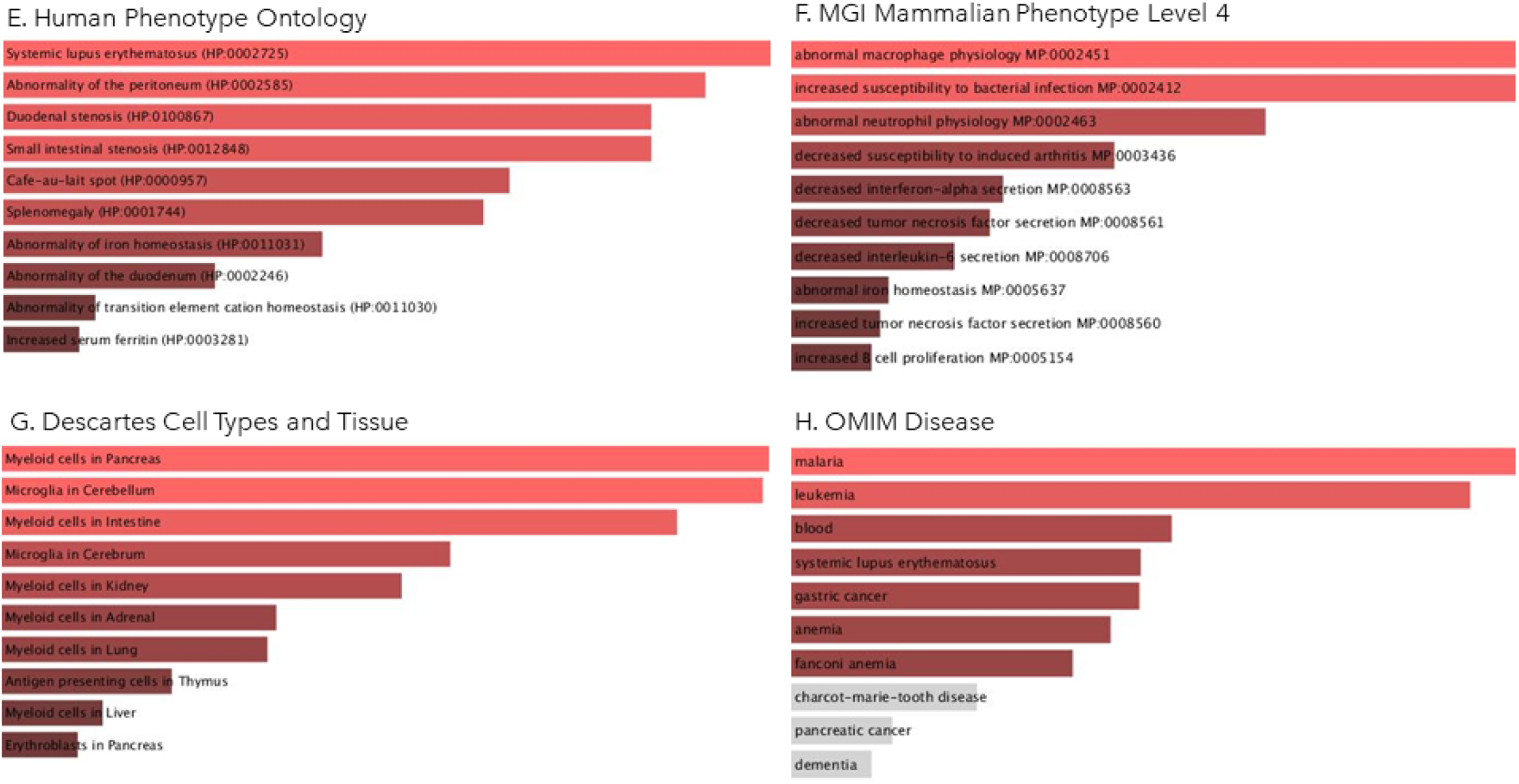
Ontologies and pathways upregulated in DESeq2 analysis of severe and asymptomatic COVID-19 subjects using Enrichr database.

**Figure 6:**
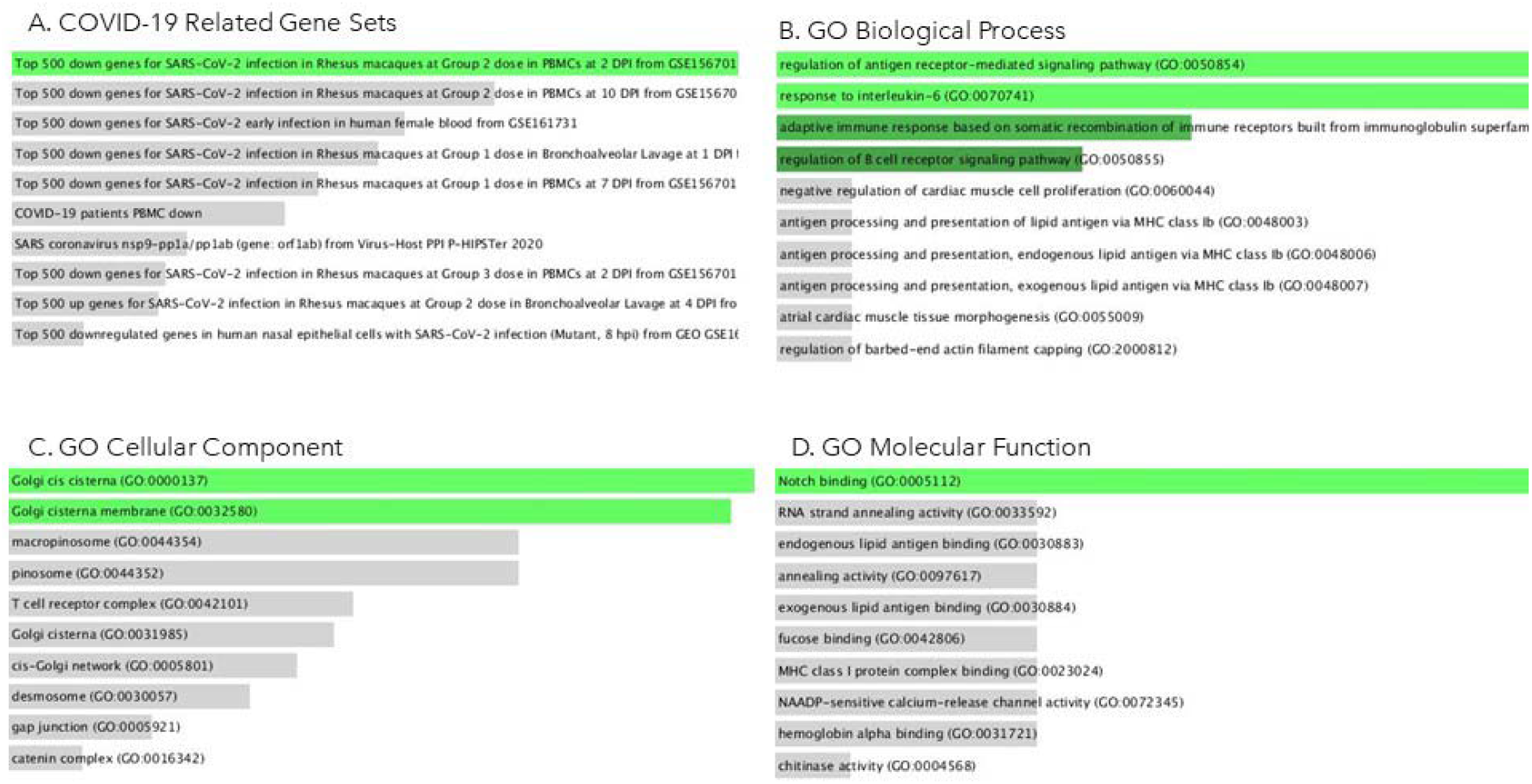
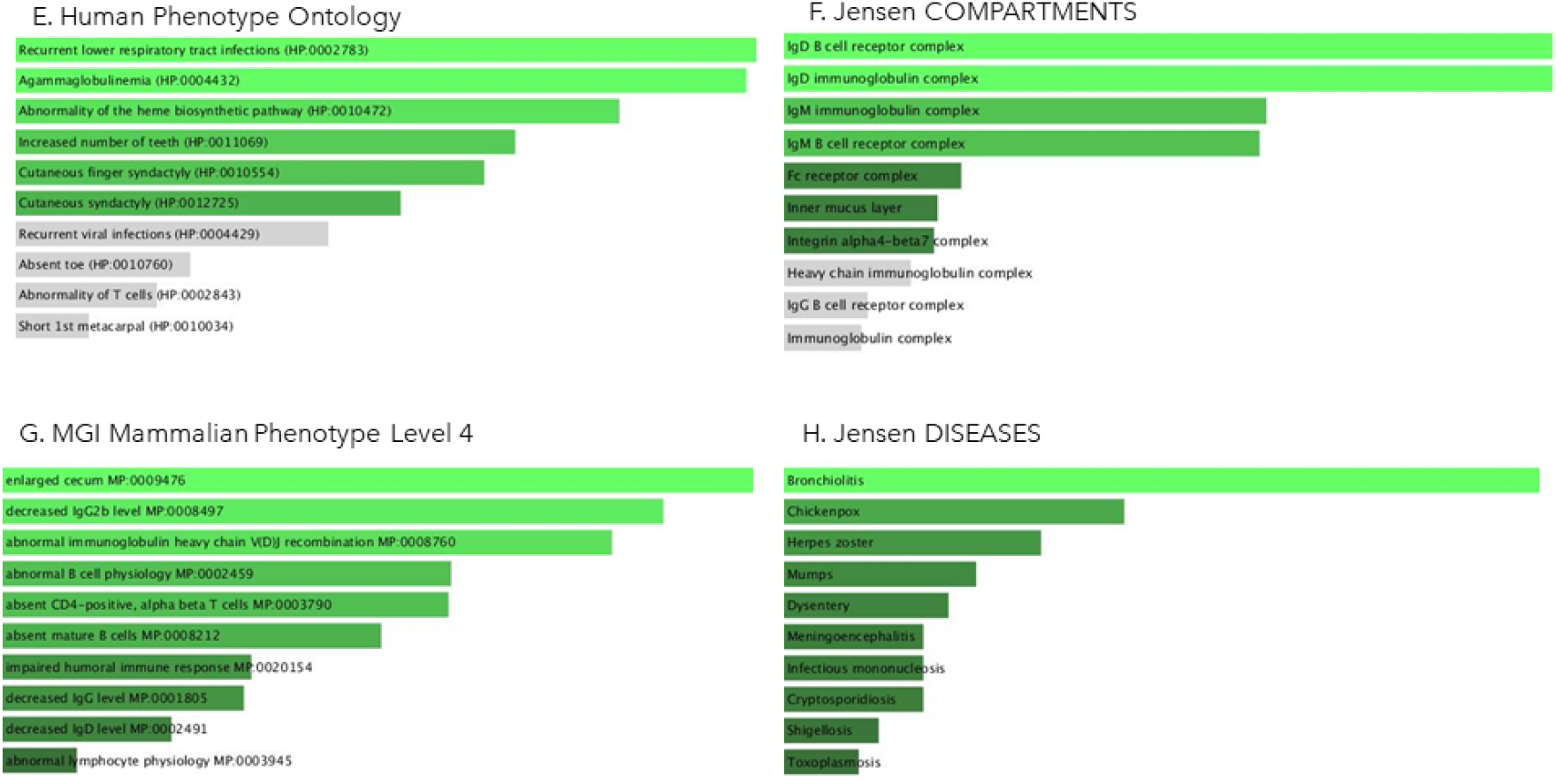
Ontologies and pathways downregulated in DESeq2 analysis of severe and asymptomatic COVID-19 subjects using Enrichr database.

##### 3.2.5.1. Gene Set Enrichment Analysis of upregulated Genes

###### Association with the viral infection and inflammatory response

Immune response terms that were found to be enriched for upregulated gene set include decreased interleukin-12b secretion MP:0008670; decreased B cell proliferation MP:0005093; abnormal interleukin level MP:0008751; impaired natural killer cell-mediated cytotoxicity MP:0005070; increased prostaglandin level MP:0009814; lymph node hyperplasia MP:0008102, Oncostatin M Signalling Pathway WP2374. Further, enriched terms found to associated with response to viral infection like Type II interferon signaling (I.F.N.G.) (WP619), IL-18 signaling pathway (WP4754), IL8 signaling (WP4754), Structural Pathway of Interleukin 1 (IL-1) (WP2637), IL-6 signaling pathway (WP364), decreased interferon-alpha secretion (MP:0008563), IL-4 signaling pathway (WP395), decreased interleukin-1 beta secretion (MP:0008658), abnormal T-helper 2 physiology (MP:0005466), abnormal macrophage physiology (MP:0002451), abnormal T-helper 1 physiology (MP:0005465), abnormal granulocyte physiology (MP:0002462), sepsis (MP:0005044). While the enriched terms related to anti-inflammatory and immune response are Activation of NLRP3 Inflammasome by SARS-CoV-2 (WP4876), abnormal inflammatory response (MP:0001845), IL-10 Anti-inflammatory Signaling Pathway WP4495.

###### Association with secondary infections

Interestingly, we found the enrichment of upregulated genes with terms that are associated with various infections other than COVID-19. These enriched terms are Influenza A, Epstein-Barr virus infection, Kaposi sarcoma-associated herpesvirus infection, *Staphylococcus aureus* infection, Measles, Human immunodeficiency virus 1 infection, Hepatitis C, increased susceptibility to bacterial infection (MP:0002412), Recurrent gram-negative bacterial infections (HP:0005420), increased susceptibility to fungal infection (MP:0005399), increased susceptibility to bacterial infection (MP:0002412), increased susceptibility to Picornaviridae infection (MP:0020937), Kaposi sarcoma-associated herpesvirus infection, increased susceptibility to Riboviria infection (MP:0020913), increased susceptibility to Herpesvirales infection (MP:0020916). Enrichment terms related to nutrients for upregulated gene set were Copper homeostasis WP3286, Vitamin B12 Disorders WP4271, and Zinc homeostasis WP3529. Iron homeostasis enrichment terms in upregulated gene sets are Ferroptosis WP4313, Folate Metabolism WP176, abnormal iron homeostasis MP:0005637, decreased spleen iron level MP:0008808, Abnormality of iron homeostasis (HP:0011031).

###### Association with organs other than the respiratory system

We also observed enriched pathways related to various organs, such as kidney-related glomerulonephritis MP:0002743; renal glomerular immunoglobulin deposits MP:0020519; and liver-related increased liver iron level MP:0008807. Heart-related Adrenergic signaling in cardiomyocytes (KEGG), myocarditis MP:0001856, Extracellular vesicles in the crosstalk of cardiac cells WP4300, ApoE, and miR-146 in inflammation and atherosclerosis WP3926, arrhythmogenic right ventricular dysplasia (Diseases) [implication of *JUP* gene in ARVD], Arrhythmogenic right ventricular cardiomyopathy (KEGG), cholesterol level (OMIM Diseases) [implication of *VNN1* gene], myocardial infarction (OMIM Diseases) [implication of PSMA6 gene], cardiomyopathy, (OMIM Diseases) [implication of *MYBPC3* gene], Chronic obstructive pulmonary disease (HP:0006510), Abnormality of lateral ventricle (HP:0030047), Abnormality of the carotid arteries (HP:0005344); Arteriovenous malformation (HP:0100026); Arterial thrombosis (HP:0004420). Pathways enriched for the intestine are “Duodenal and small intestinal stenosis,” “abnormal gut flora balance” MP:0010377, and those related to the skin were hypopigmented skin patches (HP:0001053), Urticaria (HP:0001025), Recurrent skin infections (HP:0001581), Hyper melanotic macule (HP:0001034), Recurrent bacterial skin infections (HP:0005406), Eczematoid dermatitis (HP:0000976) skin hemorrhage MP:0011514. Brain related neurological and behavioral pathways found were Inappropriate behavior (HP:0000719), Personality changes (HP:0000751), Diminished motivation (HP:0000745), Dementia (HP:0000726), Memory impairment (HP:0002354), Restlessness (HP:0000711), Vertigo (HP:0002321), Neuroinflammation WP4919, Galanin receptor pathway WP4970, Meningitis (HP:0001287).

###### Association with male infertility

Enrichment analysis of upregulated genes set shown the association with the male infertility WP4673, Abnormality of the preputium (HP:0100587), and Erectile abnormalities (HP:0100639).

###### Association with other important pathways for understanding host response

Further, we found the enrichment of upregulated genes in Ferritin, an inflammatory marker used in COVID-19 prognosis, Transcriptional cascade regulating adipogenesis WP4211, Fibrin Complement Receptor 3 Signalling Pathway WP4136. Other WikiPathway that are observed to be significantly upregulated in severe patients are IL1 and megakaryocytes in obesity (WP2865); Adipogenesis (WP236); Non-genomic actions of 1,25 dihydroxy vitamin D3 (WP4341); Vitamin D Receptor Pathway (WP2877); Myometrial relaxation and contraction pathways (WP289); Extracellular vesicles in the crosstalk of cardiac cells (WP4300). Pathways enriched related to blood cells are thrombocytopenia MP:0003179; abnormal myelopoiesis MP:0001601; impaired hematopoiesis MP:0001606; increased spleen weight MP:0004952. Descartes_Cell_Tissue_2021 shows Myeloid cells, Microglia, Antigen-presenting cells in the Thymus, Erythroblasts, Megakaryocytes in the Heart, Corneal and conjunctival epithelial cells in Eye, Vascular endothelial cells enrichment. Jensen diseases database indicates the association of upregulated genes with Arthritis, Peritonitis, Vasculitis, Periodontitis, Tularemia, Lupus Erythematosus, Boutonneuse fever, Hemochromatosis. The enriched GO cellular function(s) were azurophil granule (GO:0042582); ficolin-1-rich granule (GO:0101002); platelet alpha granule (GO:0031091). KEGG Human 2021 terms enriched in upregulated gene sets are NOD-like receptor signaling pathway; Osteoclast differentiation; Legionellosis; Lipid and atherosclerosis; Staphylococcus aureus infection; Measles; C-type lectin receptor signaling pathway; TNF signaling pathway; Rheumatoid arthritis; IL-17 signaling pathway.

##### 3.2.5.2. Gene Set Enrichment Analysis of downregulated Genes

We observed that downregulated genes in severe patients are significantly associated with Hematopoietic cell lineage and Primary immunodeficiency. Besides, they were involved in lipid metabolism, adaptive immune response, translation, recurrent respiratory infections, heme biosynthetic pathways, etc.

###### Association with metabolic pathways

Notably, some of the downregulated genes were found to be associated with metabolic pathways such as Arachidonic acid metabolism, Inositol phosphate metabolism, Histidine metabolism, Glycosylphosphatidylinositol (GPI)-anchor biosynthesis, Linoleic acid metabolism, beta-Alanine metabolism, Fructose, and mannose metabolism, Glycerophospholipid metabolism, Carbohydrate digestion, and absorption, through enrichment was not significant.

###### Association with Adaptive immune Response

Next, we observed downregulated genes are significantly enriched in GO biological processes that are associated with adaptive immune response, including regulation of antigen receptor-mediated signaling pathway (GO:0050854), response to interleukin-6 (GO:0070741), adaptive immune response based on somatic recombination of immune receptors built from immunoglobulin superfamily domains (GO:0002460), regulation of B cell receptor signaling pathway (GO:0050855), regulation of antigen receptor-mediated signaling pathway (GO:0050854), response to interleukin-6 (GO:0070741), adaptive immune response based on somatic recombination of immune receptors built from immunoglobulin superfamily domains (GO:0002460), regulation of B cell receptor signaling pathway (GO:0050855).

###### Association with translation

Some of the downregulated genes, i.e., *EIF3CL, EIF4B, EIF5AL1, PASK*, were found to be involved (although not significantly enriched) in translation processes such as the formation of the translation preinitiation complex (GO:0001731), regulation of translational initiation (GO:0006446), positive regulation of translation (GO:0045727), regulation of translational elongation (GO:0006448), cytoplasmic translational initiation (GO:0002183), translation Factors WP107.

###### Association with recurrent respiratory diseases and abnormal Heme biosynthesis

Recurrent lower respiratory tract infections (HP:0002783), Agammaglobulinemia (HP:0004432), Abnormality of the heme biosynthetic pathway (HP:0010472).

###### Other signaling pathways

Further, the TGF-beta signaling pathway, Notch signaling pathway, and Ferroptosis pathways were also associated with downregulated genes. Besides, downregulated genes are related to Cutaneous finger syndactyly (HP:0010554), Cutaneous syndactyly (HP:0012725), and Increased number of teeth (HP:0011069).

## Discussion

The primary concern with highly transmissible COVID-19 disease is the lack of understanding of the disease-causing mechanisms, resulting in poor treatments and post COVID-19 complications [110]. Clinical observations and scientific studies indicate that SARS-CoV-2 infection impacts not only respiratory organs but also other organs such as the brain, heart, kidney, gastrointestinal tract, etc. [8, 111–114]. The risk factors for COVID-19 severity include pre-existing comorbidities, particular age group of subjects, demographics, gender, etc. [5, 8, 23, 115, 116]. The heterogeneous effects of the infection on various individuals pose a significant hurdle in the therapeutic management of COVID-19 patients. Thus, it is vital to delineate the molecular alterations occurring in different groups of patients based on the impact of infections. This study extensively explored the transcriptomics profiles of two contrasting groups of COVID-19 patients, i.e., severe, and asymptomatic. The RNA sequencing data is derived from whole blood cells, a pool of immune cells, and significant biochemical products result of biochemical processes, making it a considerable tissue sample for transcriptomic profiling. Hence, we believe that the whole blood serves as a good source for understanding the immunopathology of COVID-19 subjects. Identifying significant pathways involved in the patients might help manage the severity of the disease. Exploratory analysis using Principal Component Analysis (PCA) shows distinct asymptomatic and severe subjects clusters. Subsequently, differential gene expression analysis was performed between these two groups employing the DESeq2. We identified 2,837 genes as significantly differentially expressed between severe and asymptomatic COVID-19 subjects. To reduce false discovery rate (FDR) and increase statistical significance, we used a stringent filter (Bonferroni-padj value <0.05 and Log2fold Change (Log2FC)) and found 1,224 upregulated (Log2FC >= 1.5 and p-adjusted value <0.05) and 268 downregulated (Log2FC <= −1.5 and p-adjusted value <0.05) genes in severe in comparison to asymptomatic COVID-19 subjects. Further, to understand the alterations at the molecular and biological level, we queried differentially regulated genes in the Enrichr database.

Our study found pathways known to express in viral response, in general, and specific to COVID-19 infection. We observed the type II interferon (IFNG) pathway upregulated in severe subjects. While type I IFNG is generally activated in viral response, studies have found suppression of Type I IFN in SARS-CoV infections [117, 118]. The enrichment studies observed an increased population of myeloid cells (in the pancreas, intestine, kidney, lung, liver) and microglia (in the brain), which form part of the innate immune response against the virus. These cell types have a known role in phagocytosis and anti-inflammation, biochemical pathways commonly observed in response to viral infection [119, 120]. Antigen-presenting cells (A.P.C.) in thymus enriched in severe patients also indicate an immune response to the virus. G.O. cellular components show increased ficolin and azurophil-rich granules secretion in severe subjects. These are also associated with the COVID-19 immune response [121–123]. Neutrophil count increases in COVID-19 infection [115, 124]. Our enrichment analysis also found neutrophil activation and neutrophil mediated immunity. Neutrophil degranulation is enhanced in response to inflammatory reactions in the body [125–127]. And previous studies have also reported increased neutrophil degranulation in response to COVID-19 infection in organisms other than humans [125]. One such study performed on the Rhesus macaque model shows increased neutrophil degranulation in young subjects compared to old subjects [125]. As observed in severe patients, we propose that the upregulation of neutrophil degranulation occurs in response to the disease severity. Our subjects fall in the mean age of around 45 years, the old age group; we need to compare neutrophil degranulation in COVID-19 response in an age-dependent study.

Further, our analysis observed “negative regulation of viral process” in the upregulated G.O. biological process, possibly explaining that the increased host immune response (anti-viral) reduces other viruses’ multiplication. The possible reason for this is the activation of the anti-viral immune response that reduces the risks of other infections. Increased STING (stimulator of interferon genes), which mediates interferon expression, is a known prognosis of COVID-19 [128]. Interestingly, we also observed that COVID-19 severe patients might have an increased risk of bacterial, fungal infection compared to asymptomatic patients. This observation of reduced viral infection but increased secondary infection aligns with previous studies on COVID-19 patients [129, 130]. The immune response involved in viral and bacterial infection shares different immune components [131–136]. A study reveals simultaneous expression of both IFNα and IFNγ inhibits the expression of biomarkers associated with viral and bacterial infection [131]. We believe that the complex interplay of viral and bacterial response factors and activation of viral response in the host inhibits the expression of host immune machinery to tackle bacterial infection and might be the probable reason for increased susceptibility to bacterial infections post COVID-19 infection. Previous research shows that NOD-like receptor signaling enhanced in response to SARS-CoV-1 infection results in disturbances in microbiota and increased secondary infection [118, 137]. We also observed NOD-like receptor signaling enrichment in the upregulated gene set, which might indicate gastrointestinal manifestations and increased susceptibility to bacterial infection in severe patients. This observation needs to be further confirmed by studying the host response expression induced by infections with various microorganisms.

Another significant observation in enriched terms for upregulated genes is the high coincidence of cardiac complications in COVID-19 patients, as evident in severe COVID-19 patients [17, 29]. Our study has observed coagulation dysfunction upregulated in severe patients, which could be the reason for the cardiac manifestation of COVID-19 infection. As mentioned previously, we have observed enrichment of Antigen-presenting cells in the thymus, and there are studies linking the thymus’ role in Arrhythmia [138–140]. We have also confirmed many “Disease-specific laboratory values” upregulated in severe patients. These are related to immunological response, inflammation response, and hypercoagulable state, increased aspartate aminotransferase (AST), and alanine aminotransferase (ALT), and increased interleukin 6 (IL-6), and decreased thrombocytes, reduced blood sodium.

We have also found that COVID 19 disease severity might impact fertility in the patients. It is known that ACE2 receptors are present in human male testicles [141], but the studies related to COVID-19’s impact on testicular functionality are contradictory [142–144]. Studies done to detect viral RNA in semen showed different results, with the majority indicating the absence of SARS-CoV-2 RNA [145–149]. So, we propose that if the viral particles are absent in the semen, the possibility of infertility in COVID-19 patients could be an inflammatory response to COVID-19 infection [150]. The enrichment pathways related to downregulated gene sets also support the previous research findings, such as reduced notch binding signaling, absent mature B cells, CD4+, alpha, and beta T cells, etc. [125, 151].

As COVID-19 disease has a multifactorial response on the body, we need more clinical features for prognosis, which can help us manage the diverse impact of COVID-19 on health. To address future novel virus disease management, we must not limit ourselves to real-time therapeutics. Instead, we must continuously build the concepts of generalized host response and disease progression on diverse tissues and subject groups.

## Conclusion

Our unpreparedness with SARS-CoV-2 indicates the need for more stringent research to help us understand disease progression and devise strategies for other such outbreaks in the future. Our comparative study based on two contrasting COVID-19 infection conditions, i.e. severe and asymptomatic patients identified the alteration of key pathways and biological processes associated with various comorbidities. We observed upregulation of viral-specific immune response and inflammatory pathways. Besides, heightened organ-specific responses related to blood, heart, brain, intestine, and kidney enriched in severe subjects not limited to respiratory organs. Also, our study suggests that severe COVID-19 subjects become more prone to bacterial infections and less prone to viral infections. Besides, we found the downregulation of lipid metabolism, adaptive immune response, translation, heme-biosynthetic pathways, etc. The major pathways highlighted in our study are associated with cardiac complications, autoinflammatory conditions, secondary infections, iron homeostasis and anemia, lipid metabolism, male infertility, etc. These altered pathways in severe patients might be indicative of post-COVID effects. We anticipate our study will facilitate clinicians in managing COVID-19 patients and post-COVID complications, essentially, researchers in finding better therapeutic targets. However, analyzing more samples in both groups will help validate our findings.

### Limitation of the Study

We face multiple challenges in the transcriptomic analysis of COVID-19 patients. There are limited host response data available. The lack of diversity of data (tissues, demographics, patient clinical characteristics) is also another limitation. Also, we need more data available from infection studies to establish a correlation between secondary infection and SARS-CoV-2. The study can be improved in the following ways. As observed in COVID-19 patients, the immune response components also vary during infection, and hence many of these complications may also change with the course of disease progression.

## Authors contribution

Sen P performed the data analysis, produced the figures, and wrote the manuscript. Kaur H proposed the project, reviewed the data, provided assistance with analysis, drafted the manuscript, and supervised the whole project.

## Acknowledgment

The authors are thankful to the OmicsLogic Research fellowship program, Pine Biotech, Inc, U.S.A.

## Supplementary Data

Supplementary File 1: Supplementary Figures

Supplementary File 2: Supplementary Tables

