## Supplementary File 1 for "In Silico transcriptional analysis of asymptomatic and severe COVID-19 patients reveals the susceptibility of severe patients to other comorbidities and non-viral pathological conditions"

### **In Silico transcriptional analysis of asymptomatic and severe COVID-19 patients reveals the susceptibility of severe patients to other comorbidities and non-viral pathological conditions**

Poonam Sen ^1^, Harpreet Kaur^1*^

1. Pine Biotech, New Orleans, U.S.A.

**Emails of Authors**

Poonam Sen:

Harpreet Kaur:

*** Correspondence**

Harpreet Kaur, PhD

Curriculum Developer & Research Consultant

Pine Biotech

1441 Canal St #229, New Orleans, LA 70112, United States

ORCID ID: <https://orcid.org/0000-0003-0421-8341>

Homepage: <https://www.linkedin.com/in/harpreet-kaur-phd-b06722134/>

A.
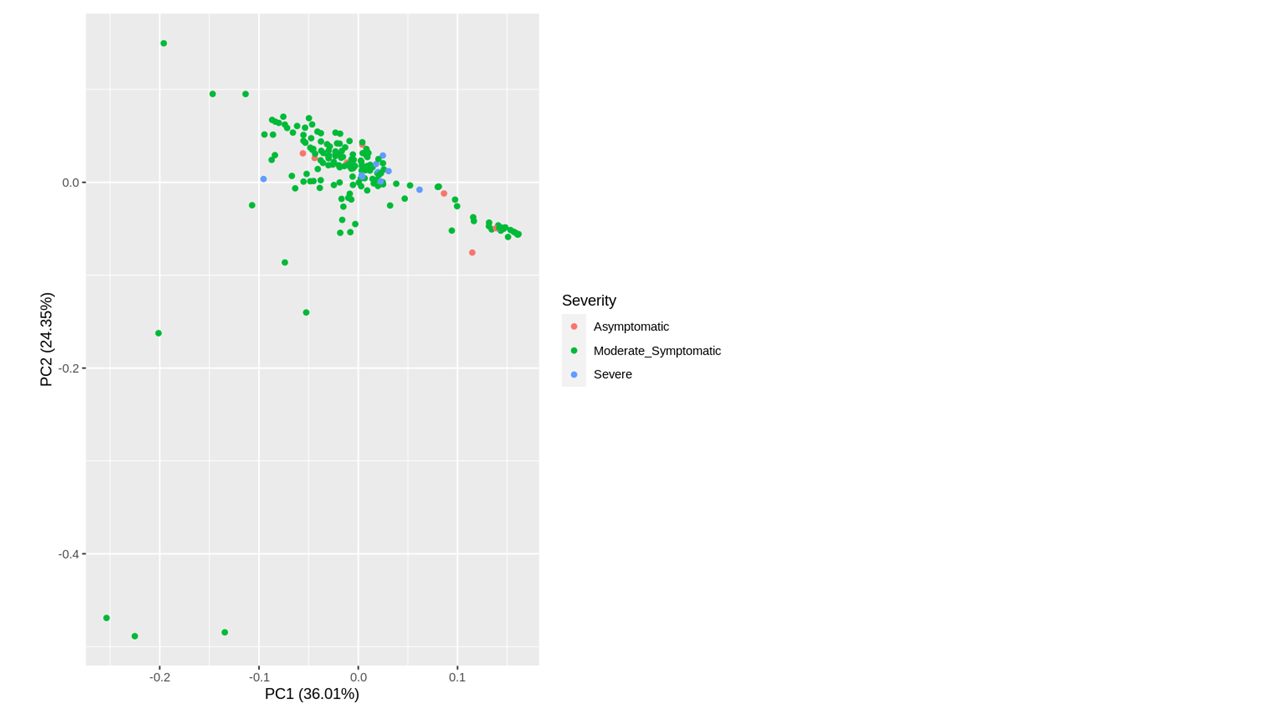

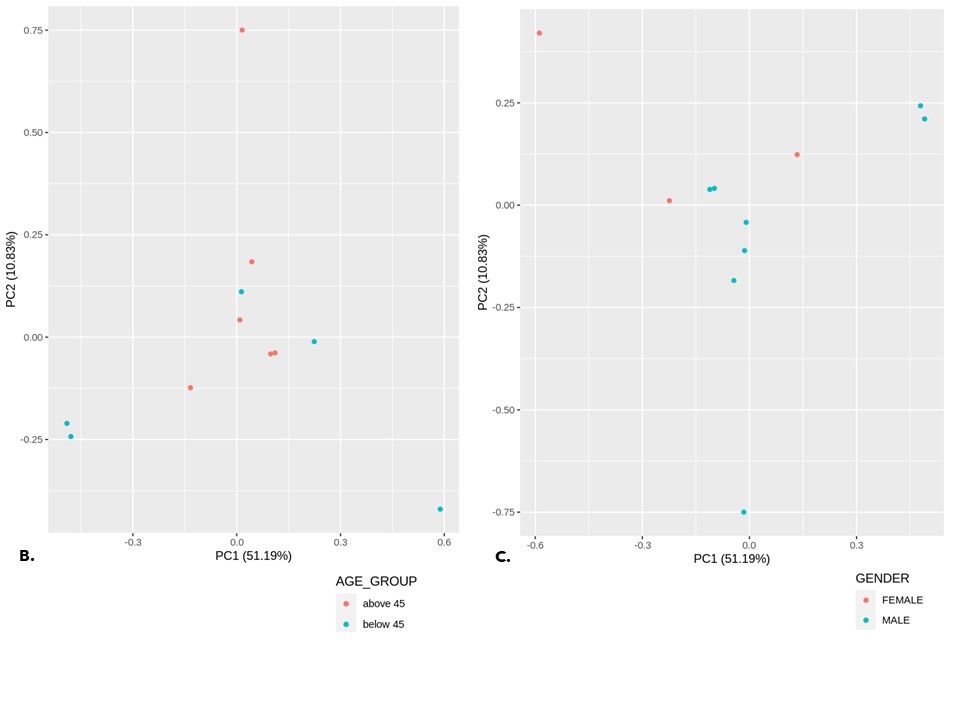
Supplementary Figure S1. Principal Component Analysis between various groups. A. The Principal Component Analysis of all 180 samples of asymptomatic, moderately symptomatic, and severe subjects (n=180, Day00 and Day05). B. PCA of Severe below 45 age (n=2, Day00) v/s Severe above 45 age (n=4, Day00) C. Severe male (n=7, Day00) v/s Severe female (n=7, Day00) C. Severe untreated (n=, Day00) v/s Severe treated (n=7, Day00)

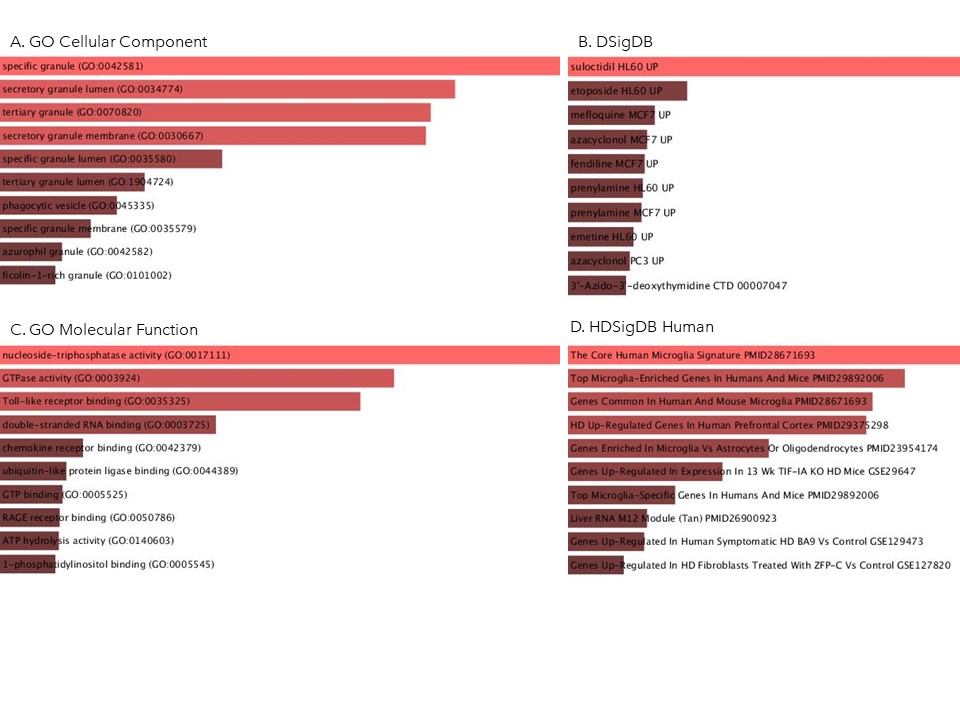

Supplementary Figure S2: Ontologies and pathways upregulated in DESeq2 analysis of untreated severe and asymptomatic COVID-19 subjects using Enrichr database. A. GO Cellular component B.DSigDB C.GO molecular function D. HDSigDB Human.

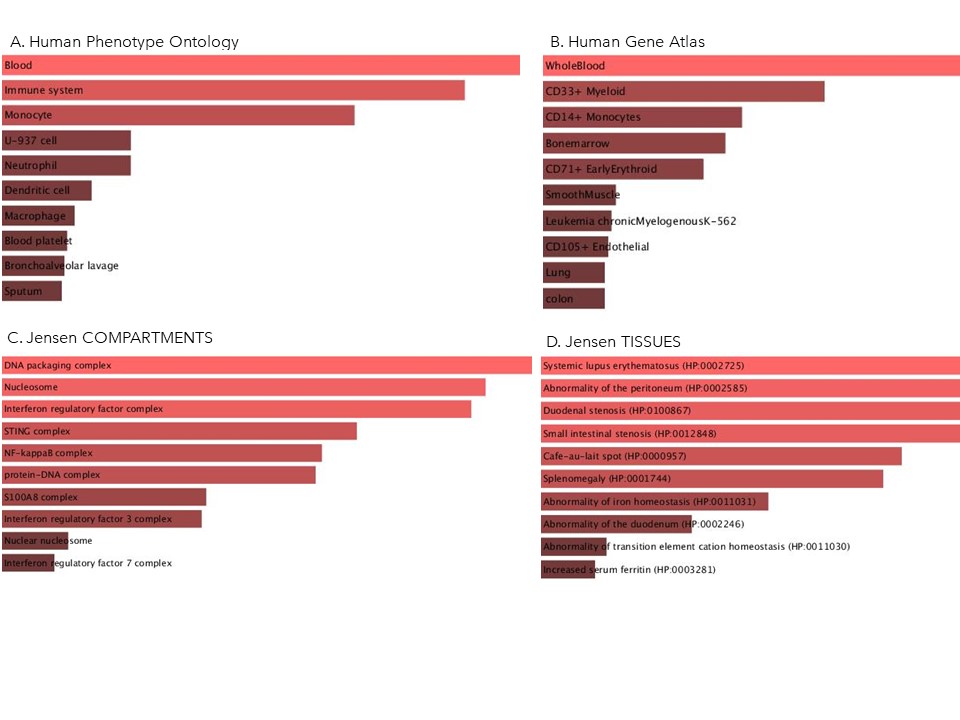

Supplementary Figure S3: Ontologies and pathways upregulated in DESeq2 analysis of untreated severe and asymptomatic COVID-19 subjects using Enrichr database. A. Human Phenotype Ontology B. Human Gene Atlas C. Jensen Compartments D. Jensen Tissues.

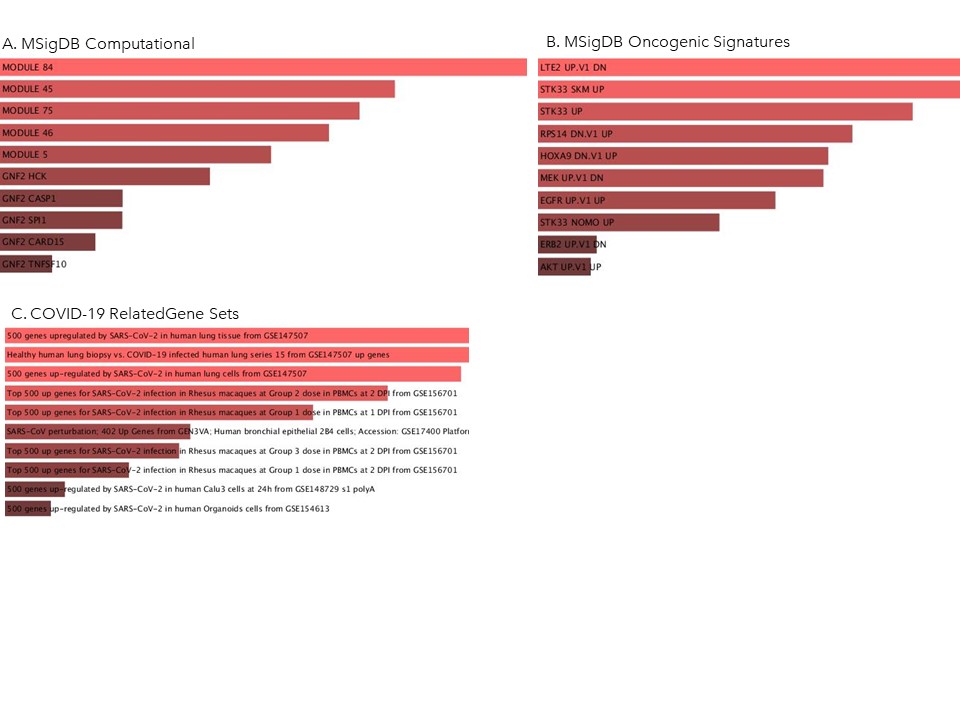

Supplementary Figure S4: Ontologies and pathways upregulated in DESeq2 analysis of untreated severe and asymptomatic COVID-19 subjects using Enrichr database. A. MSigDB Computational B.MSigDB Oncogenic Signature C. COVID-19 related gene sets.

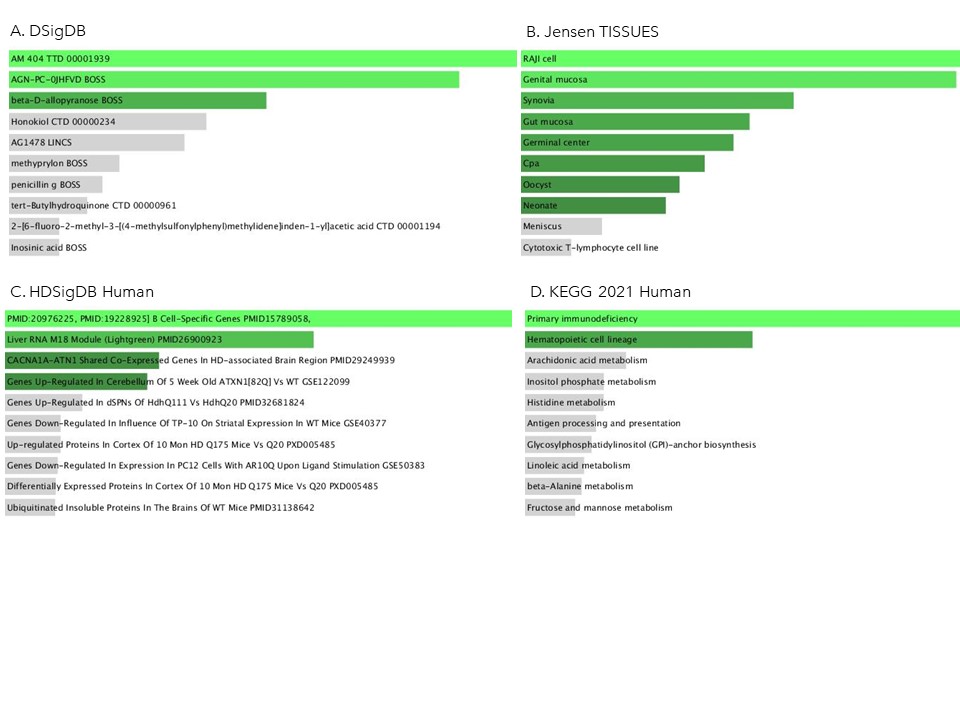

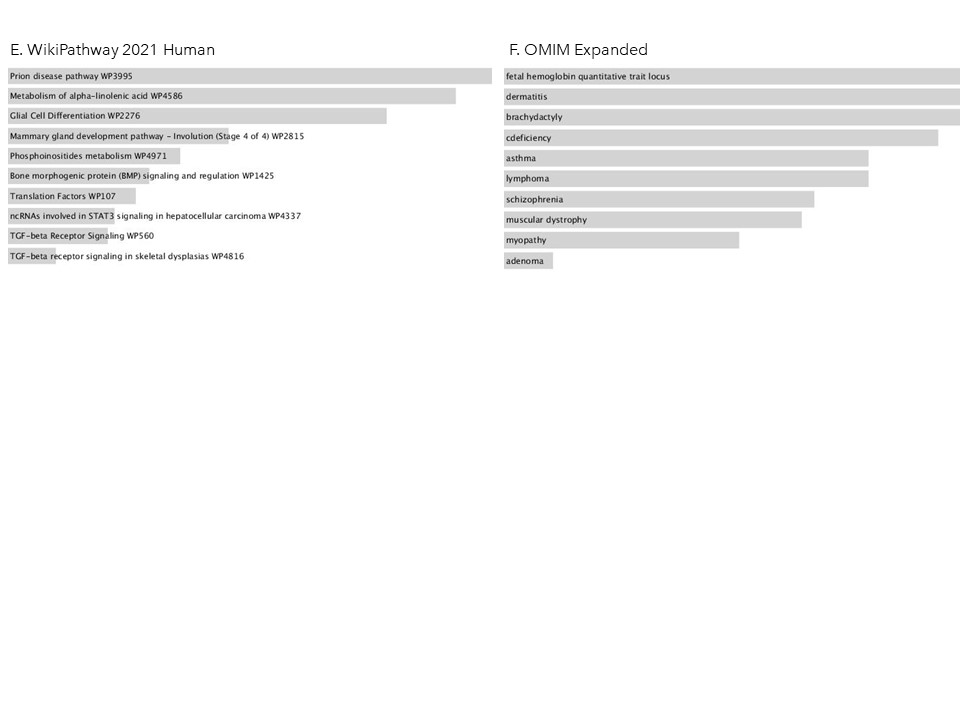

Supplementary Figure S5: Ontologies and pathways downregulated in DESeq2 analysis of untreated severe and asymptomatic COVID-19 subjects using Enrichr database. A. DSigDB. B. Jensen Tissues. C. HDSigDB Human. D. KEGG Human. E. Wikipathways Human. F. OMIM Expanded.
